# Uncovering the distinct properties of a bacterial type I-E CRISPR activation system

**DOI:** 10.1101/2021.10.06.463193

**Authors:** Maria Claudia Villegas Kcam, Annette J. Tsong, James Chappell

## Abstract

Synthetic gene regulators based upon CRISPR-Cas systems offer highly programmable technologies to control gene expression in bacteria. Bacterial CRISPR activators (CRISPRa) have been developed that use engineered type II CRISPR-dCas9 to localize transcription activation domains near promoter elements to activate transcription. However, several reports have demonstrated distance-dependent requirements and periodical activation patterns that overall limit the flexibility of these systems. Here, we demonstrate the potential of using an alternative type I-E CRISPR-Cas system to create a CRISPRa with distinct and expanded regulatory properties. We create the first bacterial CRISPRa system based upon a type I-E CRISPR-Cas, and demonstrate differences in the activation range of this system compared to type II CRISPRa systems. Furthermore, we characterize the distance-dependent activation patterns of type I-E CRISPRa to reveal a distinct and more frequent periodicity of activation.

## INTRODUCTION

The engineering of CRISPR-Cas systems has led to the development of new molecular tools for genome engineering and gene expression control. (1, 2) To date, these technologies have largely relied on the type II CRISPR-Cas9 system, which offers the convenience of a single protein effector domain and can leverage a simple, single guide RNA (sgRNA) design. To control gene expression, researchers use a catalytically inactive variant of the Cas9 protein (dCas9) that allows the complex to be targeted to DNA without cleaving it. This has been exploited to create gene regulatory systems that are facile to program to new DNA targets. In bacteria, this includes transcription repressors called CRISPR interference (CRISPRi), (3) and more recently, CRISPR activators (CRISPRa) that can turn on transcription when localized upstream of promoters. (4–9) Together these technologies allow for exciting applications such as high throughput functional genomic screening (10) and rapid strain engineering for optimizing metabolite production. (11)

Current bacterial CRISPRa systems are subject to target site requirements that limit their broad utility. Like all regulators based upon CRISPR-dCas9, CRISPRa systems require the presence of a three-nucleotide PAM (NGG) adjacent to the DNA target site. Additionally, several groups have reported other target site requirements for CRISPRa. Specifically, it has been observed that activation is only achieved when CRISPRa is targeted to PAM sites within a 2-4 bp window that repeats every 10-11 bp located upstream (~70-100 bp) of the promoter’s transcription start site (TSS). (9, 12) This is due to the requirement of the recruited activation domain (AD) to be localized on specific faces of the DNA helical plane relative to the promoter. Unfortunately, this means that for a given gene, only a fraction of target binding sites are functional, which significantly limits the overall gene targeting range.

Engineered variants of dCas9 have provided one route to expand the target range of bacterial CRISPRa systems. For example, PAM-relaxed dCas9 variants that use non-canonical PAM sites have allowed activation of promoters lacking correctly positioned NGG PAM sites. (12) Additionally, circularly permuted variants of dCas9 (cpdCas9) can allow localization of ADs to distinct positions within the tertiary structure of dCas9, and as a result, have shifted activation patterns that allow activation from PAM sites located in suboptimal positions. (9)

Exploration of different CRISPR-Cas types provide a potential alternative to create bacterial CRISPRa systems with distinct regulatory properties. For example, another CRISPR-Cas system that has been utilized for gene regulation is the type I-E CRISPR-Cas system from *E. coli*. (13) This is composed of a CRISPR associated complex known as Cascade (comprised of Cas8e, Cse2, Cas7, Cas5 and Cas6 proteins (14)), a CRISPR RNA (crRNA), and a Cas3 enzyme that performs DNA cleavage and thus is excluded for gene regulation applications. Comparing the type II and type I-E systems, there are numerous differences that we posit could give rise to distinct regulatory properties. First, the PAM site requirements of Cascade are distinct and typically more flexible than dCas9. (15) Thus, a different and broader set of PAM sites are available for a given target gene. Second, the structure of these systems present significant differences that we reason will result in distinct activation patterns. For example, the length of the bound DNA is 20 and 32 nucleotides (nt) for type II and type I-E systems respectively. More dramatic is the distinct bending of the dsDNA that results from binding. For example, DNA bending is 30o when bound to dCas9 (16) and 90o when bound to Cascade (17). We posit that these structural differences will result in distinct positioning of fused ADs relative to targeted DNA promoters, which in turn could give rise to unique activation patterns. Here we explore the potential of using type I-E CRISPR-Cas systems to create a bacterial CRISPRa and characterize its regulatory properties.

## RESULTS

### A bacterial CRISPRa design based upon the type I-E CRISPR-Cas system

We first sought to determine if the *E.coli* type I-E CRISPR-Cas system could be used to create a bacterial CRISPRa system, and to compare its activation to currently used dCas9 systems. As the basis of this platform we utilized a previously described modular CRISPRa approach that uses noncovalent protein-protein interactions called SYNZIP domains to recruit ADs to Cas. (9) The advantage of this approach is that it allows each component to be encoded on separate and interchangeable plasmid elements that are facile to exchange **(Figure 1A)**. As a point of comparison, we decided to use a CRISPRa system composed of a dCas9 with an N-terminal SYNZIP domain fusion, an AD composed of the alpha N-terminal domain (α-NTD) from *Pseudomonas aeruginosa* RNA polymerase with a C-terminal SYNZIP fusion, and a sgRNA targeting upstream of a constitutive RFP plasmid **(Figure 1B)**. The specific AD and dCas9 variant were chosen based on previously reported results showing strong activation in *E. coli* cells. (9) To create a bacterial CRISPRa using the type I-E CRISPR-Cas system, we created a plasmid encoding the Cascade complex from *E.coli* with a SYNZIP domain fused to the N-terminus of Cas8e **(Figure 1C)**, mimicking the N-terminal fusion to dCas9. We also created a crRNA expression plasmid and a genome-modified *E.coli* MG1655 strain lacking the endogenous CRISPR-Cas system (MG1655 ΔCRISPR-Cas). To measure transcription activation, we used a reporter RFP plasmid and designed a set of sgRNAs/crRNAs targeting sites located in the template strand 80 bp (−80T) or 100 bp (−100T) upstream of the reporter promoter’s TSS. *E. coli* MG1655 ΔCRISPR-Cas cells were transformed with the reporter RFP plasmid, the AD-SYNZIP plasmid, the dCas9 or Cascade plasmid, and the corresponding sgRNA/crRNA plasmid, or a no-sgRNA/-crRNA control plasmid containing only antibiotic resistance. RFP fluorescence (540 nm excitation and 600 nm emission) and optical density at 600 nm were measured for each culture **(Supplementary Figure 1)** and fold activation compared to the no-sgRNA/-crRNA control was calculated **(Figure 1D)**. From these measurements, we observed activation when the type II CRISPRa was targeted to −80T but not −100T, in agreement with previously reported results. (9, 12) Interestingly, we observed the opposite trend for type I-E CRISPRa, with strong activation observed when targeting −100T but not −80T. Additional controls were evaluated to confirm that activation only happened in the presence of all the CRISPRa elements (**Supplementary Figure 2**). These results demonstrate that the targeting range of type I-E and type II CRISPRa systems are distinct, and hint that more significant differences in activation patterns might be observed.

**Figure 1.**
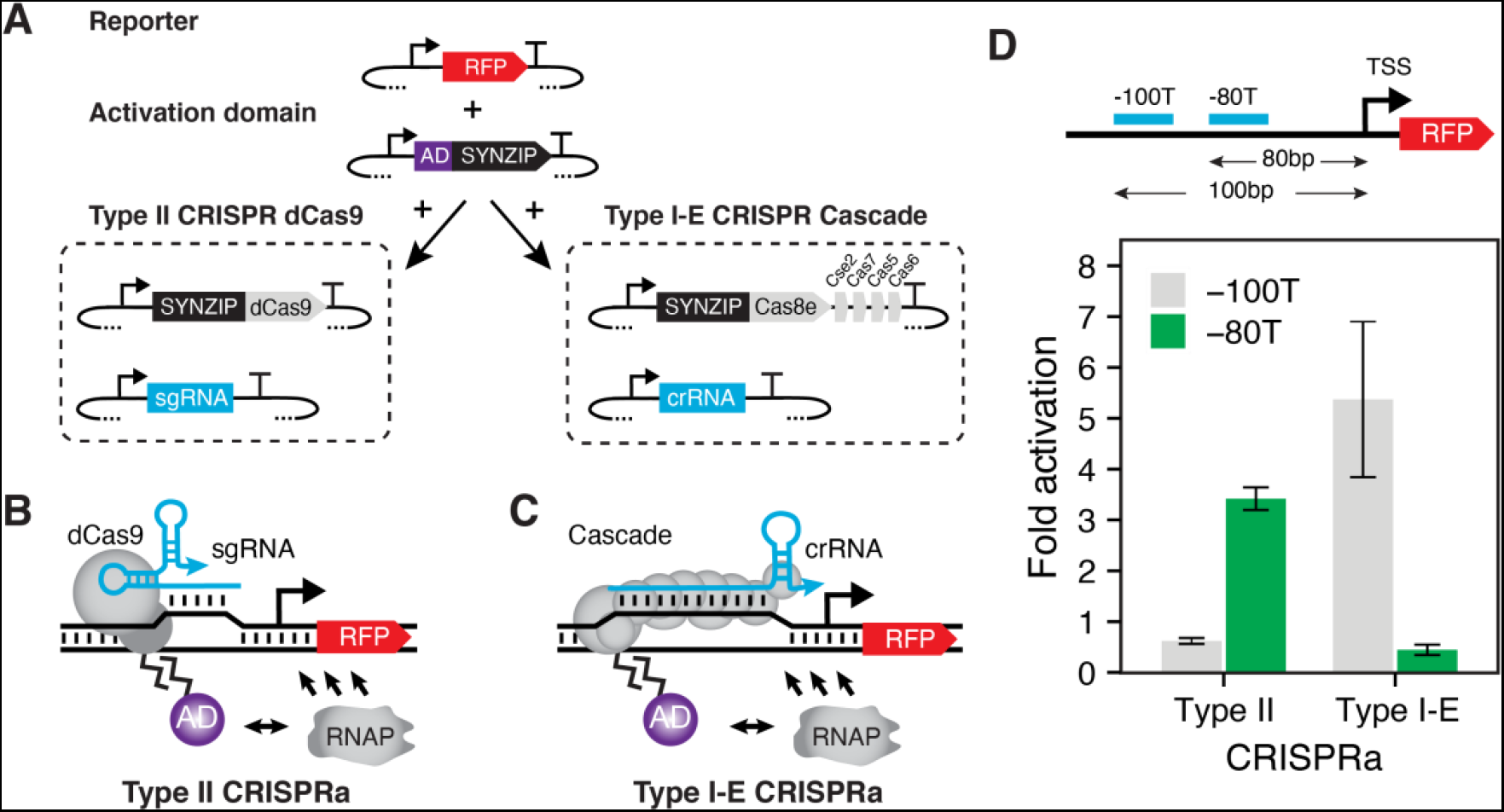
Using a modular platform to create type II and type I-E CRISPRa systems. (A)Interchangeable plasmid-encoded components of the type II CRISPRa system: RFP reporter, AD-SYNZIP, SYNZIP-dCas9, and sgRNA. For the type I-E CRISPRa system, the SYNZIP-dCas9 and sgRNA are exchanged for SYNZIP-Cascade and crRNA. Schematic of type II (B) and type I-E CRISPRa (C) that activate transcription through RNAP recruitment when targeted upstream of a target gene’s promoter. (D) Schematic of crRNA/sgRNA target binding sites and fold activation values obtained from different CRISPRa systems targeting 80 (−80T) or 100 (−100T) nt upstream of the promoter’s transcription start site (TSS). Fluorescence characterization (measured in units of fluorescence [FL]/optical density [OD] at 600 nm) was performed with *E. coli* MG1655ΔCRISPR-Cas cells transformed with plasmid combinations shown in (A). Fold activation was calculated by dividing the [FL/OD] obtained in the presence of a crRNA/sgRNA with the [FL/OD] of the no-crRNA and -sgRNA control. Data represent mean values and error bars represent s.d. of n = 4 biological replicates.

### Uncovering the distance-dependent activation patterns of a type I-E CRISPRa system

We next aimed to uncover the distance-dependent activation patterns of our type I-E CRISPRa system and to determine if these are different from the 10 bp activation patterns reported for type II CRISPRa systems. First, to get a general sense of the targeting range we identified different target sites found upstream of the reporter promoter, focusing on the template strand. Specifically, crRNAs were designed to target sites located at 71, 100, 122, 131, 142, 160 and 233 bp upstream of the TSS in the template strand of the reporter **(Figure 2A)**. We found that 2 of the 7 crRNAs located at −100T and −122T resulted in strong activation with our type I-E CRISPRa system **(Figure 2B)**. Interestingly, these activating positions are outside of the typical activation range of type II CRISPRa systems (~70 to 100 bp). (9, 12) Next, to further characterize the activation pattern, we decided to evaluate activation at 1 nt resolution between 97 and 111 bp upstream of the TSS. To do this, we adapted a previously reported approach in which a panel of crRNAs and base-shifted reporter plasmids are used to measure activation patterns at 1 nt resolution (12) by inserting additional nt between the promoter and target site **(Figure 2C)**. From this experiment, we observed a periodical activation pattern, in which strong activation is seen when the system is targeted within a 1–3 bp window that repeats every 5 bp from the TSS (**Figure 2D**). Interestingly, this pattern is distinct from the reported activation pattern for dCas9 systems that repeat every 10-11 bp. Following this, we confirmed that strong activation is seen at positions repeating every 5 bp **(Figure 2E)**. Taken together these results show a distinct set of properties for the type I-E CRISPRa system compared to type II CRISPRa.

**Figure 2.**
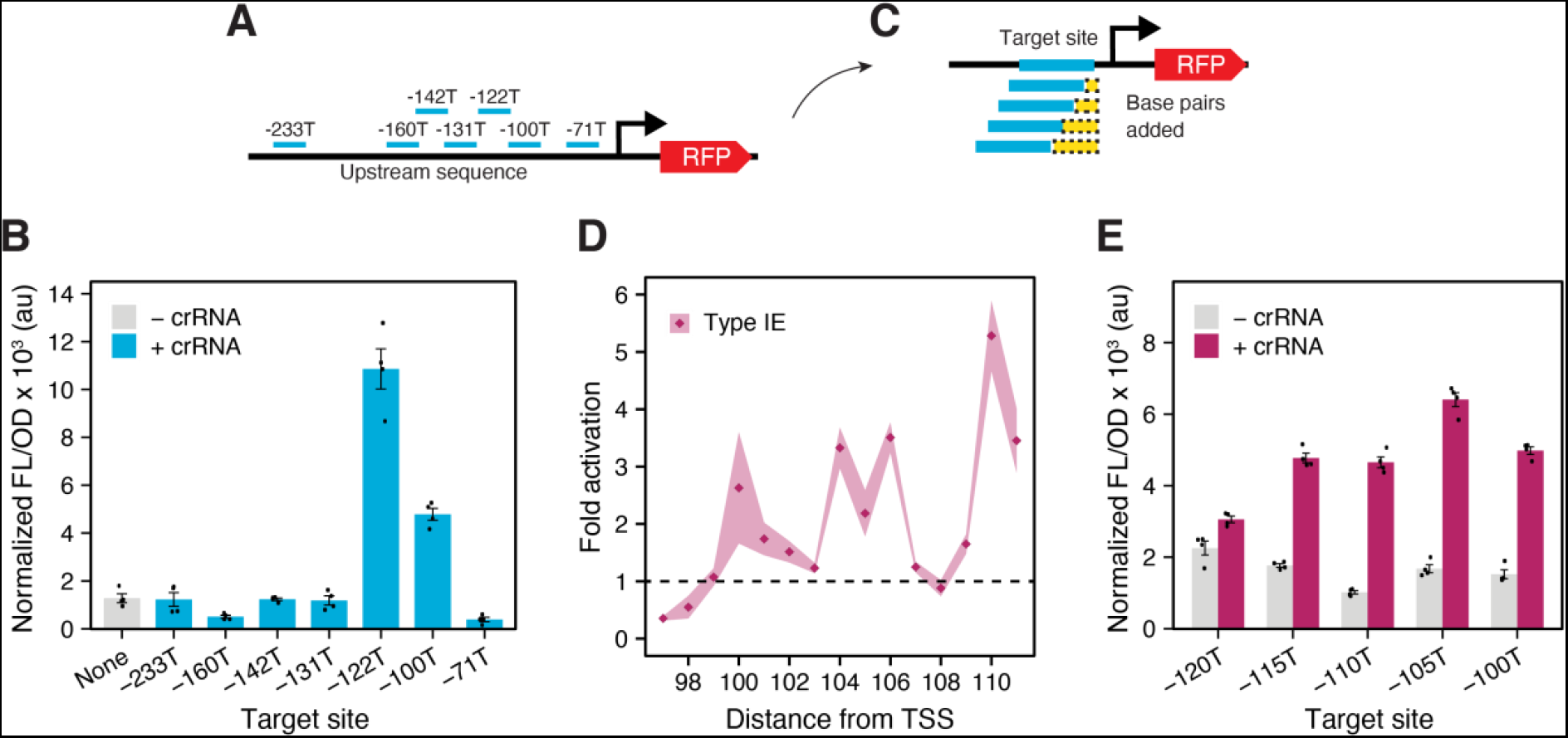
Characterization of the activation patterns for type I-E CRISPRa. (A) Schematic of the RFP reporter plasmid and crRNA target sites. (B) Fluorescence characterization of type I-E CRISPRa with crRNAs (+crRNA) targeting 7 different sites upstream of the RFP reporter gene or in the absence of a crRNA (-crRNA). (C) Schematic of the base-shifted reporter plasmid used to characterize activation patterns at nucleotide (nt) resolution. Reporters are created by inserting base pairs (yellow) between the promoter TSS and the upstream sequence, to shift crRNA target sites. (D) Fold activation showing the distance-dependent activation pattern of type I-E CRISPRa between 97 and 111 nt upstream of the reporter promoter’s TSS. (E) Fluorescence characterization of type I-E CRISPRa with crRNAs (+crRNA) targeting every 5^th^ nt between 100 and 120 nt upstream of the TSS or in the absence of a crRNA (-crRNA). Fluorescence characterization (measured in units of fluorescence [FL]/optical density [OD] at 600 nm) was performed with *E. coli* MG1655ΔCRISPR-Cas cells transformed with an RFP reporter plasmid, an AD-SYNZIP plasmid, a SYNZIP-Cascade and a crRNA-encoding plasmid or a no-crRNA control plasmid. Fold activation was calculated by dividing the [FL/OD] obtained in the presence of a crRNA/sgRNA with the [FL/OD] of the no-crRNA and -sgRNA control. Data represent mean values and error bars or shading represent s.d. of n = 4 biological replicates.

## DISCUSSION

In this work we report a novel strategy to expand the targeting range of bacterial CRISPRa systems through leveraging an alternative, type I-E CRISPR-Cas. While type I-E CRISPRa systems have been utilized for gene activators in plants (18) and human cells, (19) this is the first report of their use in bacteria. The unique activation properties observed with the type I-E CRISPRa system include a targeting distance requirement of at least 100 bp from the TSS and a 5 bp periodical activation pattern that we reason could be explained through spatial positioning of the AD relative to the promoter **(Supplementary Figure 3)**. Additionally, this work highlights the benefit of the modular CRISPRa design strategy (9) that allows for easy exchange of the CRISPR-Cas components. Given the vast repertoire and natural diversity of CRISPR-Cas systems being uncovered across the genomes of bacteria and phage, (20) we posit that this is just the beginning of exploring alternative CRISPR-Cas systems to generate and improve available technologies for gene regulation.

## ACKNOWLEDGEMENTS

The authors would like to acknowledge B. Liu for providing the MG1655 ΔCRISPR-Cas cell line. This material is based upon work supported by the Welch Foundation [C-1982-20190330 to J.C] and the Alfred P. Sloan Research Fellowship [FG-2018-10500 to J.C]. J.C. is an Alfred P. Sloan Research Fellow. The authors declare no competing financial interest.

## AUTHOR CONTRIBUTIONS

M.C.V.K and J.C. designed the study, M.C.V.K and A.J.T. performed experiments. All authors contributed to data analysis and preparation of the manuscript.

## REFERENCES

1. Xu,X. and Qi,L.S. (2019) A CRISPR–dCas Toolbox for Genetic Engineering and Synthetic Biology. Journal of Molecular Biology, 431, 34–47.

2. Ameruoso,A., Gambill,L., Liu,B., Villegas Kcam,M.C. and Chappell,J. (2019) Brave new ‘RNA’ world—advances in RNA tools and their application for understanding and engineering biological systems. Current Opinion in Systems Biology, 14, 32–40.

3. Qi,L.S., Larson,M.H., Gilbert,L.A., Doudna,J.A., Weissman,J.S., Arkin,A.P. and Lim,W.A. (2013) Repurposing CRISPR as an RNA-Guided Platform for Sequence-Specific Control of Gene Expression. Cell, 152, 1173–1183.

4. Bikard,D., Jiang,W., Samai,P., Hochschild,A., Zhang,F. and Marraffini,L.A. (2013) Programmable repression and activation of bacterial gene expression using an engineered CRISPR-Cas system. Nucleic Acids Res, 41, 7429–7437.

5. Dong,C., Fontana,J., Patel,A., Carothers,J.M. and Zalatan,J.G. (2018) Synthetic CRISPR-Cas gene activators for transcriptional reprogramming in bacteria. Nat Commun, 9, 2489.

6. Zalatan,J.G., Lee,M.E., Almeida,R., Gilbert,L.A., Whitehead,E.H., La Russa,M., Tsai,J.C., Weissman,J.S., Dueber,J.E., Qi,L.S., et al. (2015) Engineering Complex Synthetic Transcriptional Programs with CRISPR RNA Scaffolds. Cell, 160, 339–350.

7. Liu,Y., Wan,X. and Wang,B. (2019) Engineered CRISPRa enables programmable eukaryote-like gene activation in bacteria. Nat Commun, 10, 3693.

8. Ho,H., Fang,J.R., Cheung,J. and Wang,H.H. (2020) Programmable CRISPR‐Cas transcriptional activation in bacteria. Mol Syst Biol, 16, e9427.

9. Villegas Kcam,M.C., Tsong,A.J. and Chappell,J. (2021) Rational engineering of a modular bacterial CRISPR–Cas activation platform with expanded target range. Nucleic Acids Research, 49, 4793–4802.

10. Shalem,O., Sanjana,N.E. and Zhang,F. (2015) High-throughput functional genomics using CRISPR-Cas9. Nat Rev Genet, 16, 299–311.

11. Huang,C.-H., Shen,C.R., Li,H., Sung,L.-Y., Wu,M.-Y. and Hu,Y.-C. (2016) CRISPR interference (CRISPRi) for gene regulation and succinate production in cyanobacterium S. elongatus PCC 7942. Microbial Cell Factories, 15, 196.

12. Fontana,J., Dong,C., Kiattisewee,C., Chavali,V.P., Tickman,B.I., Carothers,J.M. and Zalatan,J.G. (2020) Effective CRISPRa-mediated control of gene expression in bacteria must overcome strict target site requirements. Nat Commun, 11, 1618.

13. Luo,M.L., Mullis,A.S., Leenay,R.T. and Beisel,C.L. (2015) Repurposing endogenous type I CRISPR-Cas systems for programmable gene repression. Nucleic Acids Res, 43, 674–681.

14. Xue,C. and Sashital,D.G. (2019) Mechanisms of type I-E and I-F CRISPR-Cas systems in Enterobacteriaceae. EcoSal Plus, 8, 10.1128/ecosalplus.ESP-0008-2018.

15. Xu Hua Fu,B., Wainberg,M., Kundaje,A. and Fire,A.Z. (2017) High-Throughput Characterization of Cascade type I-E CRISPR Guide Efficacy Reveals Unexpected PAM Diversity and Target Sequence Preferences. Genetics, 206, 1727–1738.

16. Jiang,F., Taylor,D.W., Chen,J.S., Kornfeld,J.E., Zhou,K., Thompson,A.J., Nogales,E. and Doudna,J.A. (2016) Structures of a CRISPR-Cas9 R-loop complex primed for DNA cleavage. Science, 351, 867–871.

17. Xiao,Y., Luo,M., Hayes,R.P., Kim,J., Ng,S., Ding,F., Liao,M. and Ke,A. (2017) Structure Basis for Directional R-loop Formation and Substrate Handover Mechanisms in Type I CRISPR-Cas System. Cell, 170, 48–60.e11.

18. Young,J.K., Gasior,S.L., Jones,S., Wang,L., Navarro,P., Vickroy,B. and Barrangou,R. (2019) The repurposing of type I-E CRISPR-Cascade for gene activation in plants. Commun Biol, 2, 1–7.

19. Pickar-Oliver,A., Black,J.B., Lewis,M.M., Mutchnick,K.J., Klann,T.S., Gilcrest,K.A., Sitton,M.J., Nelson,C.E., Barrera,A., Bartelt,L.C., et al. (2019) Targeted transcriptional modulation with type I CRISPR–Cas systems in human cells. Nat Biotechnol, 37, 1493–1501.

20. Burstein,D., Harrington,L.B., Strutt,S.C., Probst,A.J., Anantharaman,K., Thomas,B.C., Doudna,J.A. and Banfield,J.F. (2017) New CRISPR–Cas systems from uncultivated microbes. Nature, 542, 237–241.

