## Supplementary information for "Uncovering the distinct properties of a bacterial type I-E CRISPR activation system"

**TABLE OF CONTENTS:**

| | |
|---|---|
| **Page 2** | Materials and methods |
| **Page 3** | Supplementary Table 1. List of all plasmids used in this study |
| **Page 4** | Supplementary Table 2. crRNA target sequences used in this study |
| **Page 4** | Supplementary Table 3. SYNZIP-Cascade and crRNA plasmid sequences |
| **Page 5** | Supplementary Figure 1. Characterization of Type II and Type I-E CRISPRa |
| **Page 5** | Supplementary Figure 2. Controls to confirm activation with type I-E |
| **Page 6** | Supplementary Figure 3. Structure and schematics of type I-E CRISPRa activation |
| **Page 7** | References |

### MATERIALS AND METHODS

**Plasmid assembly.** All plasmids used are indicated in **Supplementary Table 1**. Most plasmids were obtained from a previously published study (1). To construct the SYNZIP-Cascade plasmid (pJEC619), Cascade genes were amplified from the genome of *E. coli* MG1655 and cloned downstream of the SYNZIP domain from the modular CRISPRa system through Golden Gate cloning. crRNA target sequences (**Supplementary Table 2**) were identified by searching the reporter plasmid (pJEC581) for available type I-E PAM sites 1-200 bp upstream of the reporter promoter's TSS. crRNA plasmids were cloned by inserting annealed oligonucleotides into a Golden Gate vector. Additional reporters for this study (pJEC701- pJEC707) were created by introducing base pairs between the upstream region and the promoter with inverse PCR.

**Fluorescence assays.** Competent cells were made from *E.coli* MG1655 ΔCRISPR-Cas strain that has the endogenous type I-E CRISPR-Cas system deleted from the genome. Cells were transformed with 3 plasmid combinations: Reporter plasmid, SYNZIP-Cas or SYNZIP-Cascade plasmid, and AD-SYNZIP plasmid or the corresponding blank plasmids and plated on LB+Agar (Difco) with 34 μg $mL^{-1}$ chloramphenicol, 25 μg $mL^{-1}$ kanamycin and 50 μg $mL^{-1}$ spectinomycin. Competent cells were prepared from the new colonies and transformed with sgRNA, crRNA, or blank plasmids. Final transformants containing 4 different plasmids were plated on LB+Agar (Difco) with 100 μg $mL^{-1}$ carbenicillin, 34 μg $mL^{-1}$ chloramphenicol, 25 μg $mL^{-1}$ kanamycin and 50 μg $mL^{-1}$ spectinomycin. Single colonies were picked and inoculated into 300 μL of LB with antibiotics in a 2 mL 96-well block (Costar). 96-well blocks were incubated at 37 °C in a Vortemp 56 (Labnet) incubator benchtop shaker with 1000 rpm shaking. After overnight growth, 10 μL of each well was diluted into 290 μL of LB with antibiotics and grown for 8-10 hours. 50 μL of each culture were diluted into 50 μL of phosphate buffered saline (PBS) solution in a 96-well plate. Optical density (OD) at 600 nm, and RFP fluorescence (FL) (540 nm excitation and 600 nm emission) were measured using an Infinite m1000 Pro plate reader (Tecan).

**Bulk fluorescence data analysis.** Each plate contained 4 biological replicates of every condition (single colonies), a set of blank control cells that contain blank plasmids with only antibiotic resistance, and a media control. Fluorescence and OD values were measured from all wells and the average value from the wells containing media was subtracted from each FL or OD. The ratio of FL to OD (FL/OD) was calculated for each biological replicate and the average FL/OD from blank control cells was subtracted from each FL/OD. Finally, mean FL/OD and standard deviations (s.d.) were calculated for each condition. To obtain the fold activation, final experimental FL/OD values from each condition containing a sgRNA or crRNA were divided by the value of the corresponding no- sgRNA/ - crRNA control. Error propagation was performed to obtain the s.d. values.

**Characterizing activation patterns.** To evaluate distance-dependent activation patterns, the crRNA (pJEC694) was used to target 11 different base-shifted reporter plasmids. The target site on each reporter is shifted by 1-10 nucleotides from the previous reporter to cover target sites at 1 nt resolution. As the basal expression of each reporter varied slightly, fold activation values were obtained using a no- crRNA control for each reporter.

**Supplementary table 1**. List of plasmids used in this study. Abbreviations are as follows: CamR = chloramphenicol resistance, AmpR = ampicillin resistance, SpecR = spectinomycin resistance, KanR = kanamycin resistance, UP = Upstream sequence, RFP = Red fluorescent protein, sgRNA = single guide RNA, crRNA = CRISPR RNA. Promoters: J23119, J23150, J23117, J23106 obtained from the iGEM Registry of Standard Biological Parts (parts.igem.org). Origin of replication: p15A, ColE1, CloDF, and pSC101. Terminators: TrrnB, t500.

| Plasmid | Name | Description | Figure |
|---|---|---|---|
| pJEC101 | Blank CamR | TrrnB – CamR – p15A ori | 1, 2 |
| pJEC102 | Blank AmpR | J23119 – TrrnB – ColE1 ori – AmpR | 1, 2 |
| pJEC103 | Blank SpecR | J23119 – TrrnB – SpecR – CloDF ori | 1, 2 |
| pJEC598 | Blank KanR | KanR – pSC101 ori | 1, 2 |
| pJEC581 | Reporter | KanR – UP – J23117 – RFP – TrrnB – pSC101 ori | 1, 2D |
| pJEC597 | Reporter +1 | KanR – UP+1 – J23117 – RFP – TrrnB – pSC101 ori | 2D |
| pJEC596 | Reporter +2 | KanR – UP+2 – J23117 – RFP – TrrnB – pSC101 ori | 2D |
| pJEC595 | Reporter +3 | KanR – UP+3 – J23117 – RFP – TrrnB – pSC101 ori | 1, 2 |
| pJEC594 | Reporter +4 | KanR – UP+4 – J23117 – RFP – TrrnB – pSC101 ori | 2D |
| pJEC593 | Reporter +5 | KanR – UP+5 – J23117 – RFP – TrrnB – pSC101 ori | 2D |
| pJEC592 | Reporter +6 | KanR – UP+6 – J23117 – RFP – TrrnB – pSC101 ori | 2D |
| pJEC591 | Reporter +7 | KanR – UP+7 – J23117 – RFP – TrrnB – pSC101 ori | 2D |
| pJEC590 | Reporter +8 | KanR – UP+8 – J23117 – RFP – TrrnB – pSC101 ori | 2D, 2E |
| pJEC589 | Reporter +9 | KanR – UP+9 – J23117 – RFP – TrrnB – pSC101 ori | 2D |
| pJEC701 | Reporter +10 | KanR – UP+10 – J23117 – RFP – TrrnB – pSC101 ori | 2B |
| pJEC702 | Reporter +11 | KanR – UP+11 – J23117 – RFP – TrrnB – pSC101 ori | 2B |
| pJEC703 | Reporter +12 | KanR – UP+12 – J23117 – RFP – TrrnB – pSC101 ori | 1, 2D |
| pJEC704 | Reporter +13 | KanR – UP+13 – J23117 – RFP – TrrnB – pSC101 ori | 2D, 2E |
| pJEC705 | Reporter +14 | KanR – UP+14 – J23117 – RFP – TrrnB – pSC101 ori | 2B |
| pJEC706 | Reporter +18 | KanR – UP+18 – J23117 – RFP – TrrnB – pSC101 ori | 2E |
| pJEC707 | Reporter +23 | KanR – UP+23 – J23117 – RFP – TrrnB – pSC101 ori | 2E |
| pJEC578 | SYNZIP-dCas9 | J23150 – SYNZIP18 – dCas9 – p15A ori– CamR | 1 |
| pJEC619 | SYNZIP-Cascade | J23150 – SYNZIP18 – Cas8e – Cse2 – Cas7 – Cas5 – Cas6 – p15A ori– CamR | 1, 2 |
| pJEC637 | AD-SYNZIP | J23106 – αNTD (P. aeruginosa) – SYNZIP17 – TrrnB – SpecR – CloDF ori | 1, 2 |
| pJEC567 | sgRNA -80 | J23119 – sgRNA80T – TrrnB – ColE1 ori – AmpR | 1 |
| pJEC583 | sgRNA -100 | J23119 – sgRNA100T – TrrnB – ColE1 ori – AmpR | 1 |
| pJEC694 | crRNA -71T | J23119 – crRNA71T – t500 – ColE1 ori – AmpR | 1, 2B |
| pJEC695 | crRNA -100T | J23119 – crRNA100T – t500 – ColE1 ori – AmpR | 1, 2 |
| pJEC696 | crRNA -233T | J23119 – crRNA233T – t500 – ColE1 ori – AmpR | 2B |
| pJEC697 | crRNA -160T | J23119 – crRNA160T – t500 – ColE1 ori – AmpR | 2B |
| pJEC698 | crRNA -142T | J23119 – crRNA142T – t500 – ColE1 ori – AmpR | 2B |
| pJEC699 | crRNA -131T | J23119 – crRNA131T – t500 – ColE1 ori – AmpR | 2B |
| pJEC700 | crRNA -122T | J23119 – crRNA122T – t500 – ColE1 ori – AmpR | 2B |

**Supplementary Table 2.** crRNA target sequences used in this study.

| Plasmid | PAM | Target sequence (5'→3') |
|---|---|---|
| pJEC694 | AAG | gcgtcctttgggttccaccggatacctccgga |
| pJEC695 | AGG | tatcctgcggtgtcctgcggttaccaaaggcg |
| pJEC696 | AAG | tgccacctgtggcaattccgacgtcgcctacg |
| pJEC697 | AGG | acacctttggttgccaagggtgacctatggtg |
| pJEC698 | AAG | ggtgacctatggtgaccatgggccaccacggg |
| pJEC699 | ATG | gtgaccatgggccaccacgggcgacctcaggt |
| pJEC700 | ATG | ggccaccacgggcgacctcaggtatcctgcgg |

**Supplementary Table 3.** SYNZIP-Cascade and crRNA plasmid sequences.

| Parts | Plasmid sequence |
|---|---|
| **SYNZIP-Cascade** (pJEC619)<br><br>CamR- Promoter J23150 - SYNZIP18 - *Cas8e* - *Cse2* - Cas7 - Cas5 - *Cas6* – dblTerm - P15A ori | [illegible] |
| **crRNA** (pJEC583)<br><br>AmpR – Promoter J23119 – Repeat1 – *32nt* – Repeat 2 – t500 – ColE1 ori | [illegible] |

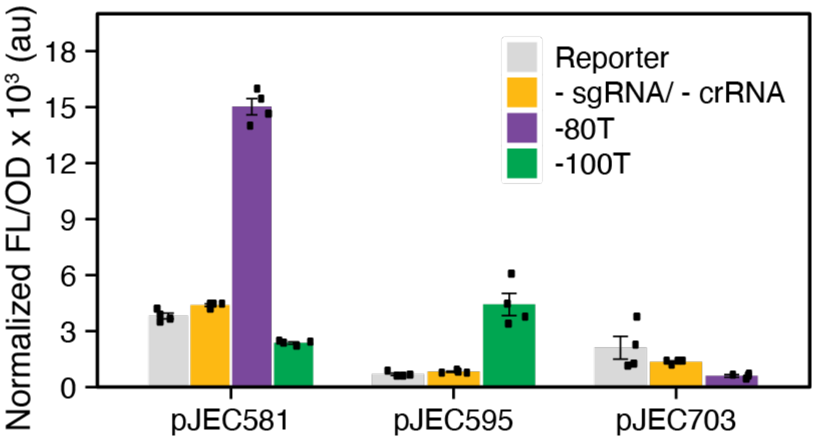


**Supplementary Figure 1. Characterization of type II and type I-E CRISPRa**. Fluorescence characterization of type II CRISPRa targeting reporter plasmid (pJEC581) at positions -80T and -100T. Fluorescence characterization of type I-E CRISPRa targeting position -100T in pJEC595 reporter plasmid and -80T in pJEC703 reporter plasmid. All three reporters express an RFP under the control of a J23107 promoter in a SC101 plasmid. Fluorescence from the reporter expressed alone is shown in grey (Reporter). Fluorescence characterization (measured in units of fluorescence [FL]/optical density [OD] at 600 nm) was performed with *E. coli* MG1655ΔCRISPR-Cas cells. Data represent mean values and error bars represent s.d. of n = 4 biological replicates.

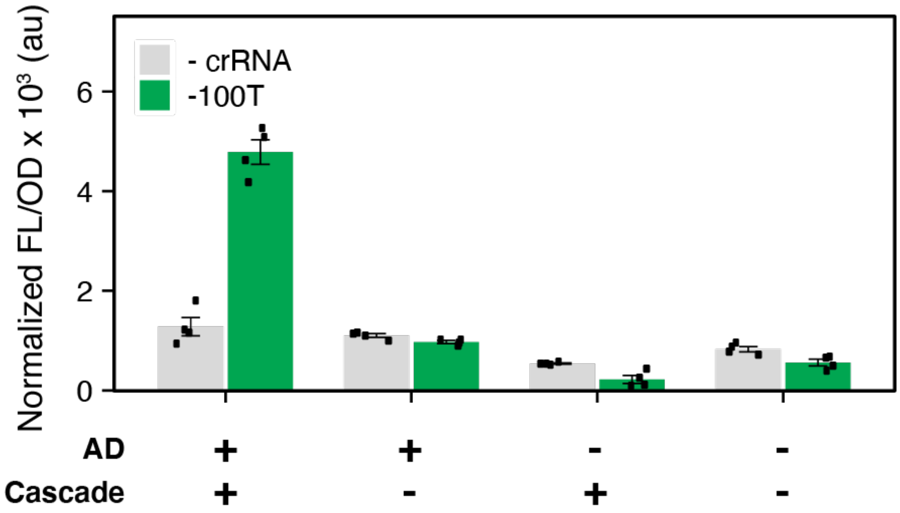


**Supplementary Figure 2. Controls to confirm activation with type I-E.** Additional controls to confirm that activation with type I-E CRISPRa targeting position -100T. Fluorescence characterization was performed in the presence and absence of each element of the CRISPRa system (crRNA, AD-SYNZIP, SYNZIP-Cascade). We show that in the absence of any one of the elements, activation is not achieved. Fluorescence characterization (measured in units of fluorescence [FL]/optical density [OD] at 600 nm) was performed with *E. coli* MG1655ΔCRISPR-Cas cells. Data represent mean values and error bars represent s.d. of n = 4 biological replicates.

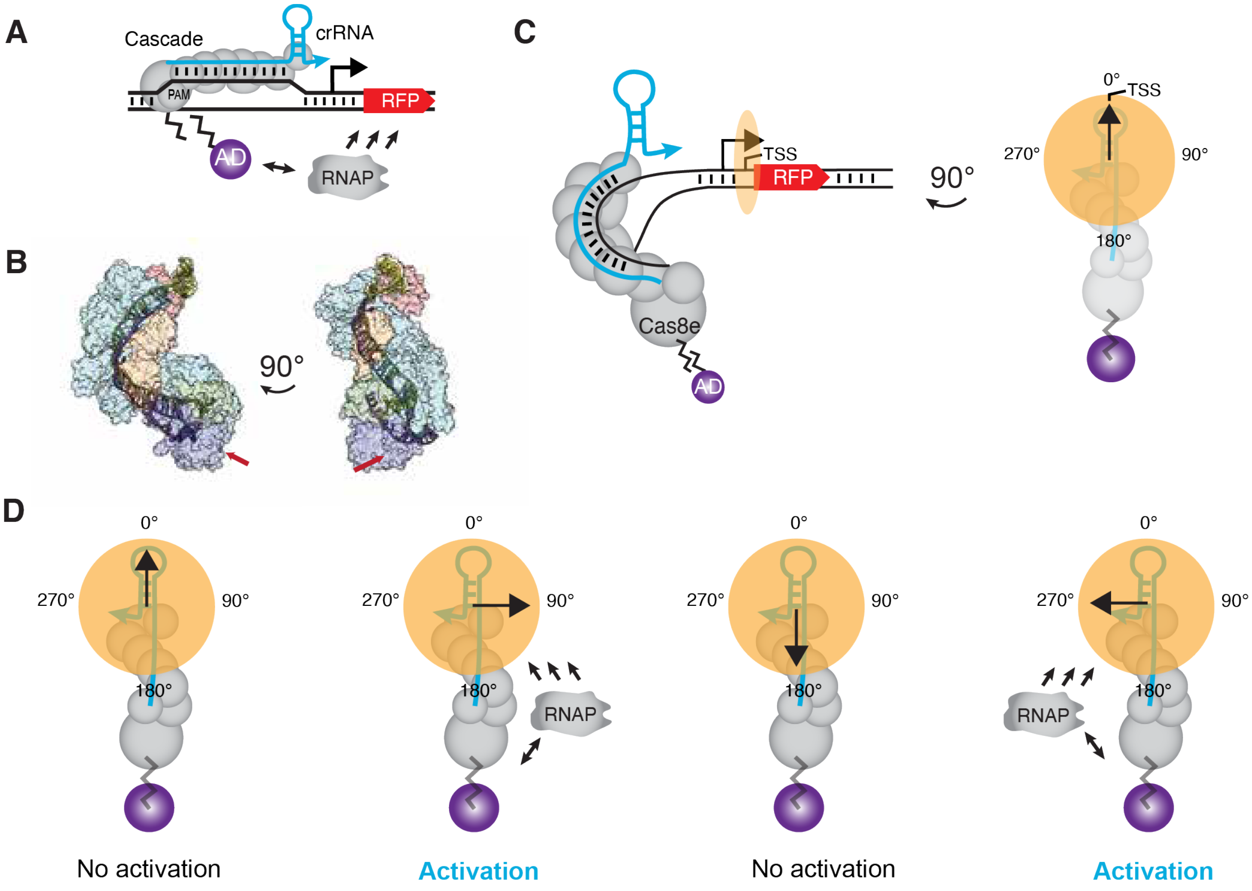


**Supplementary Figure 3. Structure and schematics of type I-E CRISPRa activation.** (A) Schematic of type I-E CRISPRa system used in this paper. (B) Crystal structure of the Cascade complex with the crRNA and DNA (PDB: 5H9E). The red arrows indicate the localization of the N-terminal fusion in Cas8e, where the AD is recruited. (C) Cartoon of the type I-E CRISPR-Cas structure from B. Orange circle used as a visual aid to indicate the position of the promoter's TSS in the helical wheel of DNA. (D) Schematic of type I-E CRISPRa recruitment of RNAP to the promoter. We hypothesize that the AD is only able to activate a targeted promoter when localized to specific surfaces of the DNA double helix (depicted as an orange circle) relative to the promoter's TSS (depicted as an arrow). One potential explanation of why activation patterns repeat with a 5 bp periodicity is if activation can be achieved on surfaces of the DNA double helix that are located ~180° apart. For example, if activation is seen when the promoter is localized at 90° or 270° within the helical wheel of DNA. Such a feature could explain why activation is seen when a crRNA is targeted to -100T and -105T, but not -102T.

References

1. Villegas Kcam,M.C., Tsong,A.J. and Chappell,J. (2021) Rational engineering of a modular bacterial CRISPR–Cas activation platform with expanded target range. *Nucleic Acids Research*, **49**, 4793–4802.
